# Scalable production of immune-silent circular RNAs for efficient protein expression

**DOI:** 10.1101/2025.11.19.689249

**Authors:** Xiaodan Liu, Huajuan Xiao, Michael Whitehead, Dhaval Varshney, Yifei Du

## Abstract

Messenger RNA-based therapeutics enable transient protein expression but are limited by its instability. Circular RNA (circRNA) offers enhanced stability, yet scalable production of high-purity circRNA remains challenging. Here, we report an oligo (dT) matrix-based purification strategy for tunable, scalable circRNA production at room temperature. By adjusting poly (A) length, circRNAs are positioned at the midpoint of the oligo (dT) elution profile, enabling one-step separation from both weakly and strongly bound RNA contaminants. We demonstrate that this approach scales linearly with comparable yields for small- and large-scale purifications. In scale-up experiments, 331 mg of highly pure circRNA was recovered in a single run. Incorporating alkaline phosphatase and RNase R digestion prior to chromatography efficiently removes immunogenic contaminants, yielding circRNAs with undetectable immune activation and robust translation in human cells. This workflow provides a practical platform for scalable production of high-quality circRNAs.

## Introduction

Delivery of synthetic messenger RNA (mRNA) into cells enables expression of encoded target genes, forming the foundation of mRNA therapeutics^1^. The success of COVID-19 mRNA vaccines has demonstrated the potential of this platform, which can also be adapted to express functional proteins for a wide range of diseases, including cancer, autoimmune disorders, genetic and metabolic diseases, and aging-related conditions^1,2^. However, the inherent instability of mRNA remains a major limitation for applications that require sustained protein expression^2^.

Circular RNA (circRNA) has emerged as a promising platform for next-generation mRNA therapeutics owing to its high stability and low immunogenicity^3-8^. CircRNAs are naturally occurring single-stranded RNA molecules with covalently closed loop structures, identified across diverse organisms from viruses to humans^3,4^. Although their biological functions remain largely unclear, recent studies suggest that circRNAs can act as sponges for microRNAs and proteins or serve as templates for protein synthesis. Translation from natural circRNAs is typically inefficient because it relies on cap-independent initiation mechanisms rather than the cap-dependent initiation used by linear mRNAs^9^. However, incorporation of viral-derived internal ribosome entry sites (IRESs) can enable efficient translation in synthetic constructs, supporting the development of circRNA-based therapeutics^8,10-12^.

Synthetic circRNAs can be generated through chemical synthesis, enzymatic ligation, or self-splicing ribozyme-based approaches^4,13,14^, with group I intron-based systems widely used for their efficiency and ability to produce long circRNAs^8,11,15-17^. Despite these advances, large-scale production remains challenging due to difficulties in achieving both high purity and scalability^4,18^. Removing impurities is critical for therapeutic applications, as even trace levels of contaminants can trigger strong immune responses^6^. Current purification strategies have notable limitations: size exclusion chromotography^11^, ion-pair reversed-phase chromatography^19,20^, and ultrafiltration^21^ offer limited resolution because of small size differences between circRNAs and their precursors; RNase R digestion is ineffective in removing highly structured RNAs^12,22^; and oligo (dT)-based methods, first used for negative selection of endogenous circRNAs^23^ and recently adapted for negative or positive selection of synthetic circRNA^17,24^, either fail to remove weakly bound species such as degraded RNA or by-products from in vitro transcription (IVT), or do not fully separate circRNAs from linear precursors. Junction-targeted circRNA capture provides high specificity but requires customized or non-reusable matrices, increasing cost and manufacturing complexity^16,25^. Therefore, a scalable, efficient, and broadly applicable purification method is urgently needed.

Here, we present a tunable selection-based purification strategy that positions circRNAs at the midpoint of the oligo (dT) binding affinity profile, enabling efficient separation from both weakly and strongly bound RNA species in a single chromatographic step. This tunable approach, combined with upstream enzymatic digestion and termed the PRO (Alkaline Phosphatase-RNase R-oligo (dT)) process, yields highly pure, non-immunogenic circRNAs with robust translational activity and scalability suitable for large-scale manufacturing.

## Results

### Oligo (dT) matrix offers tunable binding profile

The first step in our approach was to identify an RNA-binding matrix with tunable binding capacity. We selected oligo (dT) resin, a widely used and commercially available matrix for mRNA purification and manufacturing^26^. A linear nanoLuc mRNA (Lin Nluc), synthesized via IVT, was loaded onto a pre-packed oligo (dT)25 column operated on an AKTA system at 4 °C using a three-buffer system for a broad elution profile: buffer A (high salt), buffer B (no salt), and buffer C (NaOH). The Lin Nluc lacks a poly (A) tail and contains only short tandem adenine stretches (≤6 nt). As expected, Lin Nluc did not bind to the matrix and was recovered in the flow-through fraction (Figure 2A, C). When a poly (A) sequence (∼120 nt) was added using poly (A) polymerase, most of the Lin Nluc shifted to the NaOH elution, the strongest elution condition, indicating that RNAs with poly (A) lengths between 6 nt and 120 nt can be positioned at distinct regions across the elution profile (Figure 2B-C).

**Figure 1.**
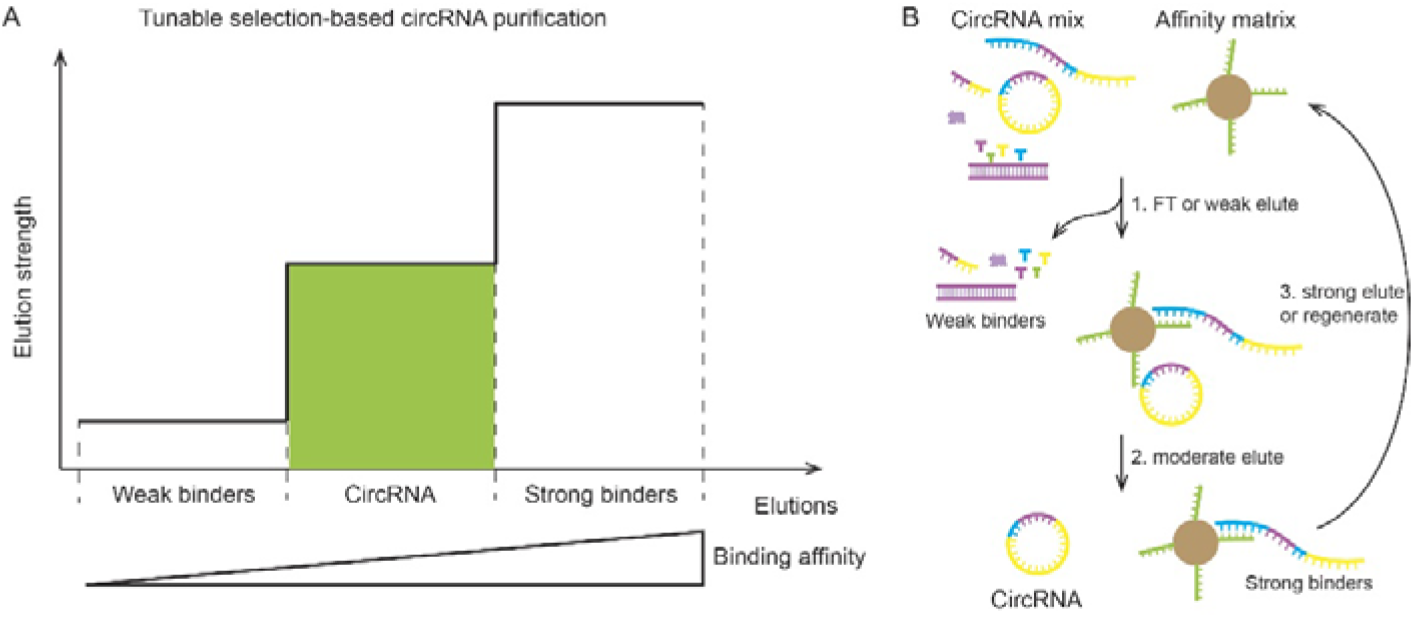
Conceptualization of the tunable selection-based circRNA purification. (A) A binding matrix captures multiple RNA species from a circRNA-containing sample, which are subsequently eluted according to their binding affinities. (B) Weaker binders, including degraded RNA, by-products of in vitro transcription (IVT), double-stranded RNA, free NTPs, and residual protein are removed in the flow-through (FT) or weak elute. The circRNA elutes under moderate conditions. Strong binders can then be eluted to regenerate the binding matrix for reuse. Unlike conventional negative or positive selection methods, this tunable selection-based approach positions circRNA at the midpoint of the elution profile (green area), enabling its efficient separation from other contaminants.

**Figure 2.**
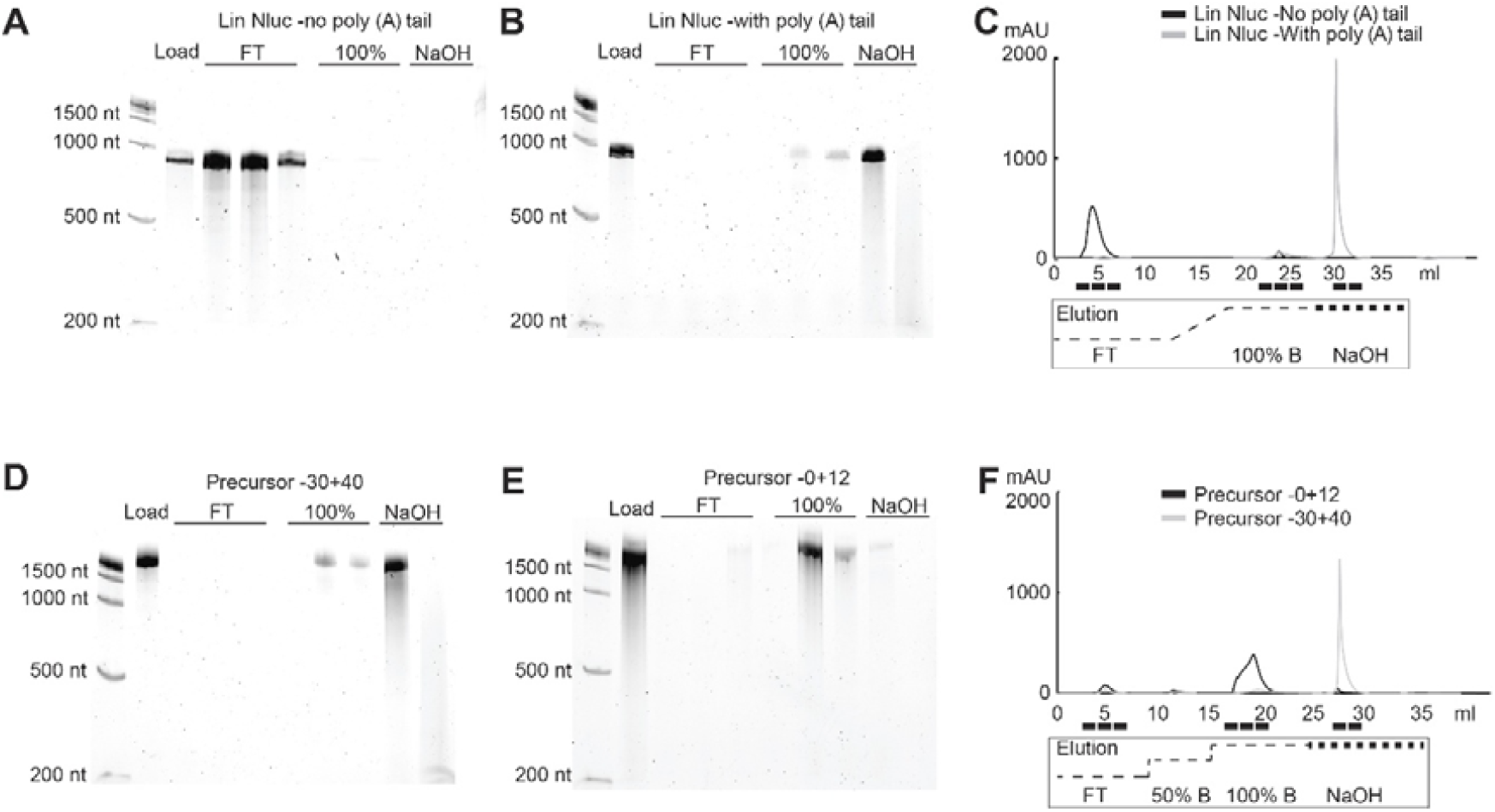
Oligo (dT)25 resin offers tunable binding capacity. Linear RNAs containing no poly (A) tailing (A), or poly (A) sequences derived either from E Coli. poly A polymerase (B) or plasmid DNA (D-E) bind differentially to the oligo dT(25) column based on their poly (A) length. Samples were loaded to an oligo (dT)25 column operated on a AKTA system at 4 °C. Elution profiles of (B) and (E) are shown in (C) and (F). FT: flow through.

We next incorporated 5′ 30 nt and 3′ 40 nt poly (A) sequences into the precursor used for circRNA synthesis based on the trans ribozyme-based circularization (TRIC) method^8^, while the circRNA region remained devoid of poly (A) sequences. The resulting elution profile showed that 30 + 40 nt poly (A) tails were sufficient to retain precursor RNAs predominantly in the NaOH fraction (Figure 2D). Reducing the poly (A) sequence to 0 + 12 nt shifted precursor RNA to 100% B fractions, a weaker elute condition (Figure 2 E-F). Together, these results demonstrate that the oligo (dT)25 matrix provides a tunable RNA binding capacity dependent on poly (A) length, establishing the basis for selective circRNA purification.

### Positioning circRNA at the centre of the Oligo (dT)25 binding profile

An effective purification process should achieve efficient separation of circRNA from both weakly and strongly bound RNA species, an outcome that previous oligo (dT)-based strategies have not achieved^17,24^. Because binding affinity to oligo (dT) resin is determined by poly (A) length, we hypothesized that designing precursor RNAs with long poly (A) sequences and circRNAs with shorter poly (A) segments would position the circRNAs at the midpoint of the elution profile, while separating it from precursor RNA that would stay strongly bound.

To test this, precursor RNAs carrying 30 + 40 nt poly (A) tails were used to generate circRNAs containing short poly (A) stretches (7–17 nt) within the circRNA region. The circularization mixture–including circRNA, linear precursors, spliced group I intron, IVT by-products, and degraded RNAs–was applied to an oligo (dT)25 column at 4 °C. CircRNAs with 7 nt poly (A) eluted primarily in the flow-through with unbound species, whereas increasing the poly (A) length gradually shifted elution toward higher elution strength (Figure 3). Because precursors with 30 + 40 nt poly (A) sequences were partially eluted at 100% buffer B (Figure 2D), we used 90% B as a moderate elution condition to achieve complete separation. Upon checking all the constructs, circRNAs containing 11-13 adenines were predominantly enriched at the midpoint of the elution profile, enabling clean resolution from both strongly bound precursors and weakly bound contaminants (Figure 3 C-D).

**Figure 3.**
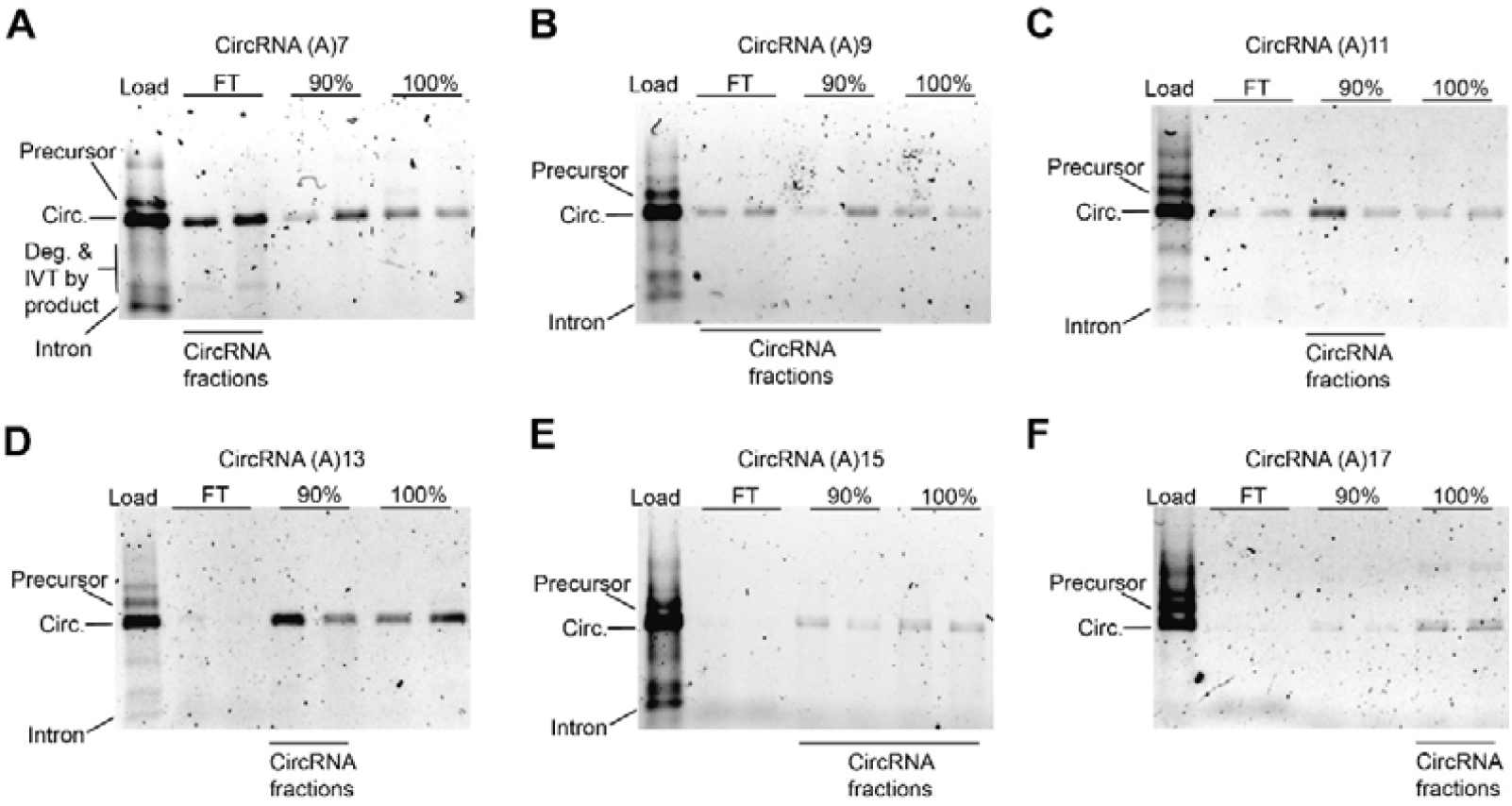
Binding of circRNAs with varying length of poly (A) sequences to oligo (dT)25 column. (A-H) CircRNAs with poly (A) sequence length ranging from 7 to 17 were tested for their binding to the oligo (dT)25 column. All precursors carry 30 + 40 nt poly (A) tails. Samples were analyzed using native agarose gels. Circ.: circRNA; Deg.: degraded RNA.

### Purification of circRNAs with poly (AC) sequences at scale

Poly (AC) and poly (A) sequences are known to enhance translation efficiency from circRNAs^11,12^. We hypothesized that these sequences alone might be sufficient to support circRNA binding to the oligo (dT)25 matrix. To test this, we examined a construct containing a poly (AC) sequence (polyAC-v1) and found that the corresponding circRNAs were strongly retained on the column (Figure 4A), preventing effective separation from precursors and the spliced group I intron, similarly to the positive selection-based purification^24^. Introducing seven A-to-C substitutions within the poly (AC) region (polyAC-v2) shifted circRNA elution to 90% buffer B, resulting in clear separation of circRNA from all other RNA species (Figure 4B–C). Interestingly, circRNA showed reduced smearing when the sample was treated with RNase R prior to loading onto the oligo (dT) column (Figure 4B).

**Figure 4.**
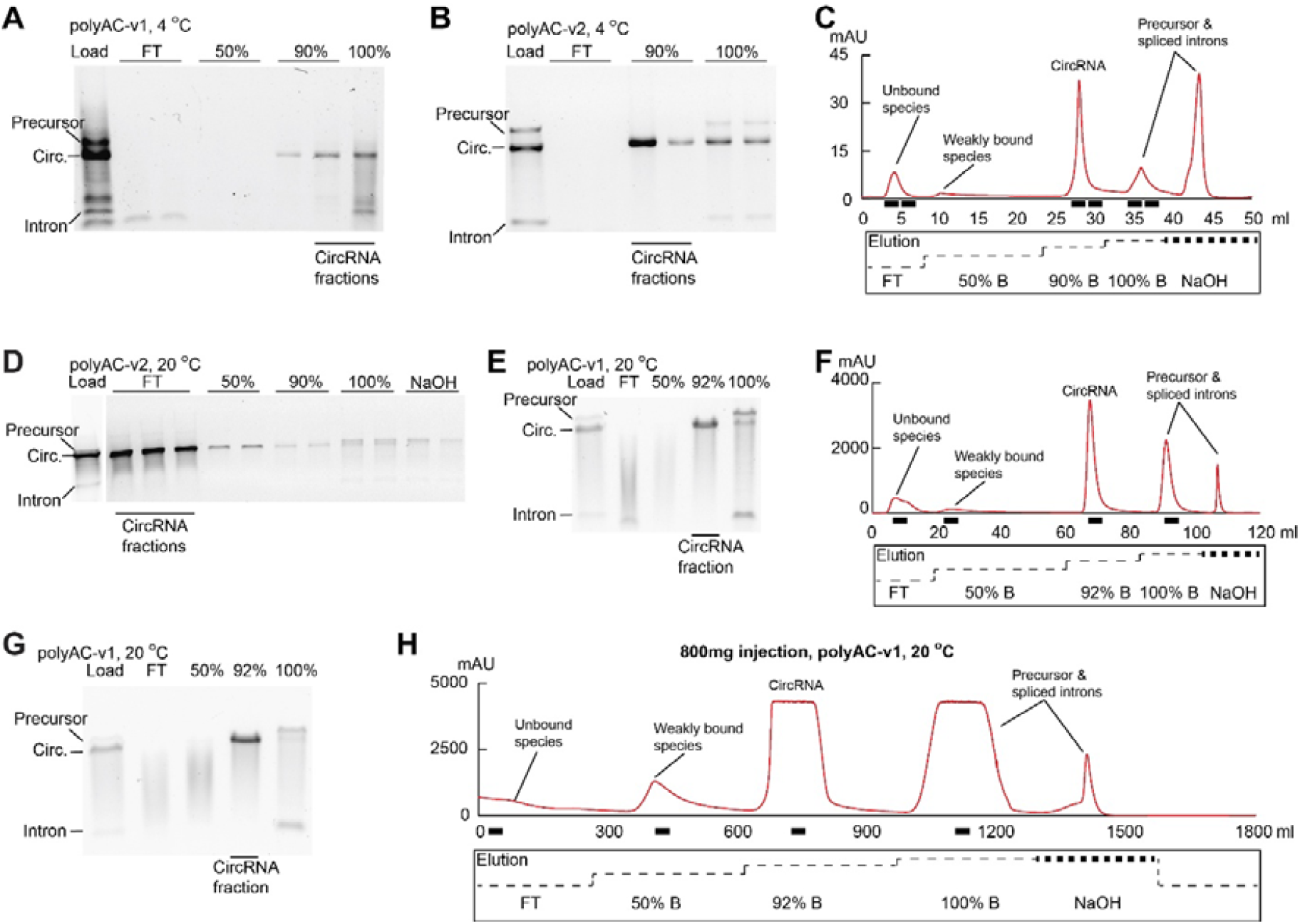
Large-scale circRNA production using oligo (dT)25 column at room temperature. (A) CircRNA containing polyAC-v1 sequences binds tightly to the oligo (dT)25 column at 4 °C. (B–C) Seven A-to-C mutations were introduced into polyAC-v1 to generate polyAC-v2; circRNA with the polyAC-v2 sequence elutes at 90% B. The circularized sample was treated with RNase R (0.5 U per μg RNA) prior to injection. (D) When purification is performed at room temperature (20 °C), circRNA with polyAC-v2 elutes in the flow-through. (E–F) Under the same conditions, circRNA with polyAC-v1 elutes at 92% B. (G–H) A single 800 mg injection of circularized RNA yielded 331 mg of highly pure circRNA.

All purifications described above were performed at 4 °C; however, maintaining this temperature complicates large-scale manufacturing. We therefore transitioned the approach to room temperature (20 °C). Interestingly, circRNAs containing the polyAC-v2 sequence were predominantly recovered in the flow-through fractions along with unbound species, similar to the negative selection-based purification^17^ (Figure 4D). In contrast, the polyAC-v1 circRNA was efficiently eluted at 90% buffer B at room temperature, suggesting that the polyAC-v1 sequence is compatible with large-scale production (Figure 4E-F). To further assess scalability, we performed large-scale purification using a 255 ml oligo(dT) column operated at room temperature. A total of 800 mg of circularized RNA mixture was loaded in a single run, yielding 331 mg of purified circRNA (Figure 4G–H). Notably, the yield from the 255 mL column was comparable to that from the 5 mL column (41.3% vs. 43.6%).

These results demonstrate that the method is reproducible and capable of large-scale circRNA production at room temperature.

### PRO purification produces immune-free circRNAs with high translation efficiency

A key advantage of circRNAs for therapeutic applications is their inherently low immunogenicity^6-8^. We previously demonstrated that a combined alkaline phosphatase (AP) digestion, RNase R digestion, and size-exclusion chromatography workflow effectively produced non-immunogenic circRNAs^8^. Building on this, we integrated the AP and RNase R digestion steps upstream of oligo (dT) purification to establish the PRO (Phosphatase-RNase R-oligo(dT)) process.

Optimization of the PRO workflow using varying enzyme concentrations showed that all tested conditions yielded high-quality circRNAs with no visible smearing or contaminant species (Figure 5A). To assess immunogenicity, mRNA levels of RIG-I and CCL5 were measured following transfection of purified circRNAs into A549 cells. Modified linear mRNA induced significantly higher RIG-I and CCL5 expression than any of the Circ Nluc RNAs (Figure 5B). In contrast, circRNAs showed no detectable immune activation under most conditions, except at the lowest AP concentration tested (0.05 U per µg RNA) (Figure 5B). We also evaluated activation of Toll-like receptors (TLRs) 3, 7, and 8, and the results indicate that PRO-purified circRNAs do not elicit a detectable TLR response (Supplementary Figure 1). Translation assays in HEK293T and A549 cells further showed that Circ Nluc treated with 1 U AP and 0.5 U RNase R per µg RNA exhibited the highest protein expression levels (Figure 5C-D). Taken together, these results demonstrate that the PRO process can produce immune-silent circRNAs with robust translation efficiency.

**Figure 5.**
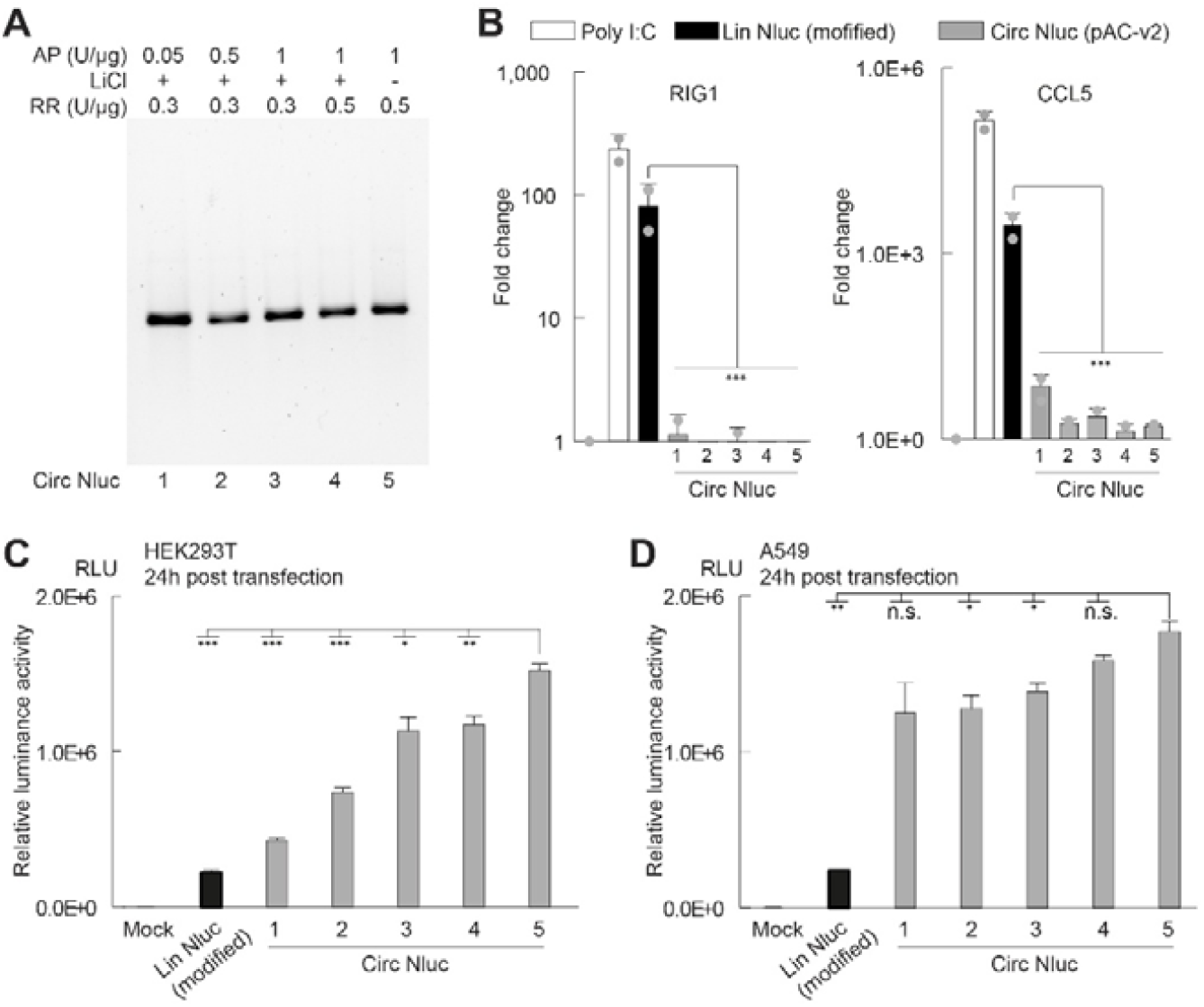
Immune-silent circRNAs production using the PRO process. (A) CircRNAs containing the polyAC-v2 sequence with 5’ 30 nt and 3’ 40 nt poly (A) sequences in the precursor were treated with varying amounts of alkaline phosphatase (AP) and RNase R (RR) prior to loading onto the oligo (dT)25 column. CircRNAs were either precipitated with LiCl2 or not precipitated before RR treatment. (B) RT-qPCR analysis of RIG-I and CCL5 mRNA levels post circRNA transfection. (C–D) Nluc expression measured in HEK293T and A549 cells. Data in (B-D) are presented as mean□±□s.d. from 3-4 biological replicates. *P□<□0.05, **P□<□0.01, ***P□<□0.001; unpaired two-tailed t-test.

### PRO purified circRNA exhibits enhanced translation efficiency

CircRNAs can also be purified using an RNase R-only approach, which simplifies scalability by eliminating chromatographic steps^12^. However, this method may not fully remove immunogenic contaminants^22^. To evaluate this, we generated circRNAs using either the PRO process or the RNase R-only protocol. As shown in Figure 6A, the RNase R-only method left residual linear precursors and spliced introns even after extended digestion, whereas the PRO workflow produced highly pure circRNA. Consistent with this observation, RNase R-only purified circRNAs induced significant immune activation and exhibited reduced protein expression in HEK293T and A549 cells compared with PRO-purified circRNAs (Figure 6B-D). These results confirm that the PRO process effectively eliminates immunogenic contaminants while yielding highly pure circRNAs with enhanced translational efficiency.

**Figure 6.**
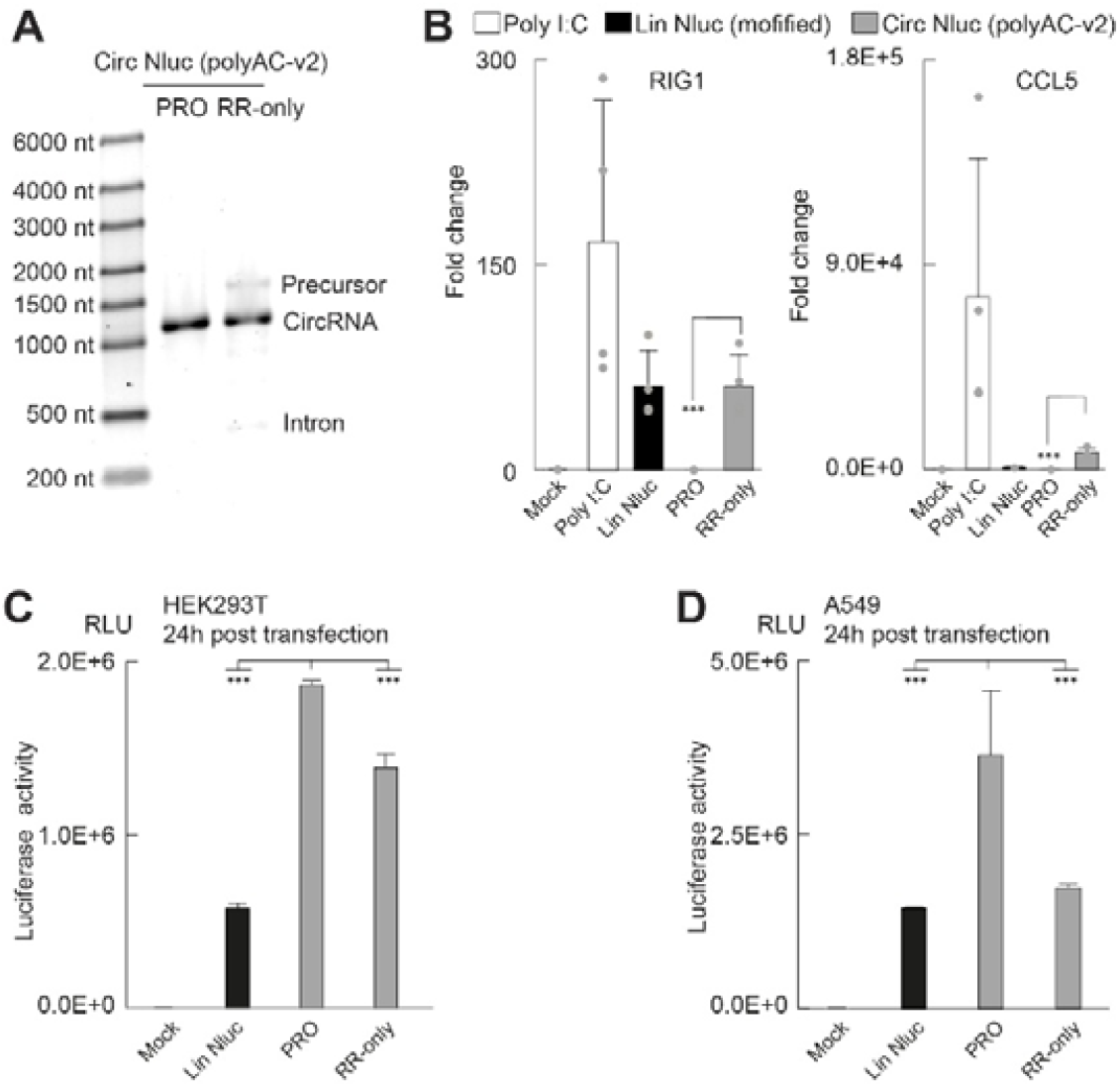
Comparison of immunogenicity and expression for PRO and RNase R-only purified circRNAs. (A-D) Gel analysis (A), immunogenicity analysis (B), and expression analysis (C-D) of circRNAs purified using the PRO or RNase R-only methods. RT-qPCR was performed to detect RIG-I and CCL5 transcript levels following transfection into A549 cells. Protein expression was evaluated in HEK293T and A549 cells. Data in (B-D) are presented as mean□±□s.d. from three biological replicates. ***P□<□0.001; unpaired two-tailed t-test.

## Discussion

Scalable production of circRNAs remains a major challenge for developing circRNA-based therapeutics^4,18^. The PRO process described here provides a simple and effective solution for purifying high-quality circRNAs with robust translational activity at scale. By tuning poly (A) lengths in circRNAs and their precursors, circRNAs were precisely positioned at the midpoint of the oligo (dT) binding profile, enabling one-step separation from both weakly and strongly retained contaminants, distinct from previously reported positive- or negative-selection-based purification strategies^17,24^. Unlike junction-specific capture approaches, PRO relies on generic, commercially available oligo (dT) resins and standard chromatography systems^16,25^, making it readily adaptable for circRNA manufacturing. The process is effective at room temperature and highly reproducible during scale-up without reduction in yield, as demonstrated by the purification of 331 mg of circRNA from 800 mg of input in a single run. Importantly, incorporation of alkaline phosphatase and RNase R digestion ensures that PRO-purified circRNAs are free of detectable immune activation and exhibit high translational efficiency in human cells.

In summary, this study demonstrates that the PRO process enables scalable, reproducible purification of immune-silent circRNAs with robust translation, providing a practical and generalizable platform for therapeutic circRNA production.

## Materials and methods

### Cloning and DNA template

The Lin Nluc and polyAC-v1 TRIC constructs were generated as previously described^8^. To introduce poly (A) sequence into the TRIC precursor, 30-nt and 40-nt poly (A) sequences were added to the 5′ and 3′ ends of the polyAC-v1 TRIC construct, respectively. The polyAC-v2 TRIC construct was generated by introducing seven A to C substitutions within the poly (AC) region. For constructs containing 7–17 nt poly (A) tracts within the circRNA region, the poly (AC) sequence upstream of the IRES was replaced with a random sequence containing the designated number of adenosines, whereas the region downstream of the stop codon was replaced with the 3’ untranslated region of human hemoglobin alpha 1 mRNA.

All DNA templates used for in vitro transcription (IVT) were cloned into a pUCIDT vector between a T7 promoter and restriction sites (NotI + EcoRV). Plasmids were propagated in homemade TOP10 competent E. coli cells and sequence-verified by Sanger sequencing (Genewiz). For IVT, plasmids were amplified in 150 mL LB cultures and purified using the Maxi Plus plasmid purification kit (QIAGEN). The purified plasmids were linearized with either EcoRV-HF (NEB), followed by deproteinization with phenol:chloroform:isoamyl alcohol (25:24:1, Sigma) extraction and isopropanol precipitation.

### RNA synthesis and circularization

RNAs were synthesized by in vitro transcription (IVT) reactions containing 35-50 μg ml^-1^ DNA template, 14 μg ml^-1^ homemade T7 RNA polymerase, 40 U ml^-1^ RNase inhibitor (Promega), 6 mM of each NTP, 1 U ml^-1^ inorganic pyrophosphatase (Thermo Fisher Scientific), and 1× IVT buffer. For the Lin Nluc, UTP was substituted with N^1^-pseudouridine triphosphate and the 1x IVT buffer contained 80 mM Tris-HCl (pH 7.4), 2 mM spermidine, 40 mM DTT, and 24 mM MgCl_2_. While for synthesis of full-length TRIC precursors, the Mg^2+^ concentration in the1x IVT buffer was reduced to 10 mM. Reactions were incubated at 37 °C for 2h for Lin Nluc and 6 h for TRIC precursors, followed by digestion with RNase-free DNase I (NEB) for 20 min. RNA was precipitated by adding an equal volume of 7.5 M lithium chloride (Sigma) and placing at −80 °C for 30 min to overnight. Precipitates were collected by centrifugation at 13,000 rpm for 20 min, washed with 75% ethanol, air-dried, and dissolved in RNase-free water.

For poly (A) tailing of Lin Nluc or full-length precursors, E. coli poly A polymerase (NEB) was used following the manufacturer’s instructions. Lin Nluc used in immunogenicity analysis and expression assay was further capped using the Vaccinia capping system (NEB) and purified by gel filtration column (Sepax, 215980P-4630). For RNA circularization, full-length precursors were first refolded by heating at 65 °C for 3 min and cooling on ice for 3 min. Refolded RNAs were then supplemented with 1/10 volume of 10× circularization buffer (500 mM Tris-HCl, pH 7.4; 100 mM MgCl_2_; 10 mM DTT; 20 mM GTP) and incubated at 55 °C for 5 min at a concentration of 500ng/ul. Reactions were terminated by adding EDTA to 10 mM, and circularized RNAs were precipitated again with 7.5 M lithium chloride, washed, and redissolved in RNase-free water for subsequent use.

### Oligo (dT) purification

When enzymatic treatments were performed prior to column injection, RNAs were sequentially treated with alkaline phosphatase (Roche) and RNase R (antibodies-online), with enzyme amounts varied to determine optimal conditions. An optional lithium chloride precipitation step was performed between these treatments. Following RNase R digestion, RNA samples were precipitated with lithium chloride (Sigma L9650), washed, and resuspended in RNase-free water.

Before column injection, RNAs were refolded by heating at 65 °C for 3□Min, followed by incubation on ice for 3□Min, and 1/10 volume of 5□M NaCl (Invitrogen) was added.

Samples were then applied to either a 5□ml prepacked oligo (dT)25 column (ThermoFisher Scientific™, A48607) or a 255□ml home-packed column (HiScale 50/20, Cytiva) containing POROS™ Oligo(dT)25 affinity resin and operated on an AKTA system (Cytiva). Flow rates were 3□ml/min for the 5□ml column and 25□ml/min for the 255□ml column.

After injection, columns were washed first with buffer A (10□mM Tris-HCl, pHL6.5, 500□mM NaCl), followed by buffer B (10□mM Tris-HCl, pHL6.5, 1□mM EDTA) applied either stepwise or as a gradient. Finally, columns were washed with 0.1□M NaOH. Fractions were analyzed using native agarose gels, and target fractions for immunogenicity and expression assays were pooled and precipitated with isopropanol.

### Cell culture and transfection

HEK293T (ECACC, Catalogue no. 12022001) cells and A549 cells (ECACC, Catalogue no. 86012804) were cultured in DMEM with 10% FBS (High Glucose GlutaMAX, Life Technologies Ltd) in a SANYO incubator at 37 °C with 5% CO_2_.

For immunogenicity analysis, A549 cells (100,000 per well) were seeded in 24-well plates (Corning) in 500 μl of medium. Each well was transfected with 200 ng of poly(I:C) (Merck) or other RNA samples using MessengerMax transfection reagent (Invitrogen). After 24 hours, cells were harvested and lysed, and total RNA was isolated using TRIzol (Invitrogen) followed by purification with the RNA Clean & Concentrator Kit (Zymo Research). The RNA was eluted in 10 μl of RNase-free water and stored at −80 °C until use in quantitative RT-PCR (RT-qPCR) analysis.

For Nluc expression, 0.04 pM of each RNA was transfected into 10,000 cells in 100 μl in 96-well plates (Corning) using the MessengerMax transfection reagent (Thermo Fisher Scientific). The Nluc assay was performed according to the manufacturer’s instructions (Promega). Briefly, HEK293T and A549 cells were lysed by adding 100□μl of lysis buffer (25□mM Tris–HCl, pH 7.4, 150□mM NaCl, 10□mM MgCl_2_, 1□mM EDTA, 2% glycerol, and 1% Triton X-100) to each well after removing the culture medium. Following a 10□Min incubation at 37□°C, 5□μl of each cell lysate was mixed with 5□μl of freshly prepared assay reagent in a black 384-well plate (Greiner). After a 3□Min incubation, luminescence was measured using a Spark 10□M plate reader (Tecan) with a 1,000□Ms integration time.

### RT-qPCR

For each sample, 300 ng of total RNA was reverse transcribed using SuperScript IV (Thermo Fisher Scientific) with a primer mixture of random hexamers and oligo(dT). Subsequently, 4 μl of each 20-fold–diluted cDNA was combined with 6 μl of reaction mix in a 384-well plate (Applied Biosystems). Each target was analyzed in triplicate. qPCR reactions were prepared using iTaq Universal SYBR Green Supermix (Bio-Rad) and primers listed in Supplementary Table 1. Amplification was performed on a ViiA 7 Real-Time PCR System (Thermo Fisher Scientific) following a standard comparative CT protocol. Data were analyzed using QuantStudio Real-Time PCR Software v1.3.

### TLR immunogenicity assay

HEK-Blue™ reporter cell lines expressing TLR3 (Invivogen hkb-htlr3) and TLR 8 (Invivogen hkb-htlr8) were cultured in DMEM, 4.5 g/l glucose, 2 mM L-glutamine, 10% (v/v) fetal bovine serum (FBS), Pen-Strep (100 U/ml-100 μg/ml), 100 µg/ml Normocin™, 30 μg/ml of blasticidin and 100 μg/ml of Zeocin®. HEK-Blue™ reporter cell lines expressing TLR7 (Invivogen hkb-htlr7v2) were cultured in DMEM, 4.5 g/l glucose, 2 mM L-glutamine, 10% (v/v) heat-inactivated FBS, 100 U/ml penicillin, 100 μg/ml streptomycin, 100 µg/ml Zeocin® and 200 µg/ml of hygromycin B. HEK-Blue™ null cells (Invivogen hkb-null1) were cultured in DMEM, 4.5 g/l glucose, 2 mM L-glutamine, 10% heat-inactivated fetal bovine serum (FBS; 30 min at 56 °C),100 μg/ml Normocin™, Pen-Strep (100 U/ml-100 μg/ml) 100 μg/ml of Zeocin®. 20ug/ml R848 provided the positive control for TLR 7 and TLR 8. Poly I:C provided positive control for TLR 3 cells. Endotoxin free water was used as a negative control. For poly I:C and RNA samples, transfection mix was made as per manufacturer’s instructions using Optimem™ and Lipofectamine MessengerMax to give final RNA concentration of 5ng/ul, 2ng/ul and 0.5ng/ul. Mock transfection mix (without RNA) was also included as a negative control.

The immunogenicity assay was performed as per manufacturer’s instructions. Briefly, 20µl of each test sample was added per well in duplicate in a 96-well plate. The growth media was discarded and the HEK-Blue™ cells were washed in warm PBS. Cell dissociation was achieved by incubating with PBS for 2 mins at 37°C. Cells were counted using Countess 3 cell counter and ∼50,000 cells in 180μl HEK-BlueTM detection medium were seeded per well containing the test samples. The plate was incubated overnight at 37 °C in 5% CO_2_ and plate was read using a plate reader at 620-655 nm.

## Statistics

All statistical analysis was done in Microsoft Excel. The Student’s t-test for two samples with unequal variances was used for significance analysis. P□<□0.05 was considered as statistically significant.

## Acknowledgments

The authors would like to thank Dr. Venki Ramakrishnan for helpful discussions and advice.

## Author contributions

Y.D. conceived the project. X.L., H.X., M.W., D.V., and Y.D. performed the experiments and analysed the data. Y.D. wrote the manuscript with inputs from all authors.

